# Fish population dynamics and diversity in boreal and temperate reservoirs: A quantitative synthesis

**DOI:** 10.1101/033282

**Authors:** Katrine Turgeon, Christopher T. Solomon, Christian Nozais, Irene Gregory-Eaves

**Affiliations:** McGill University, Department of Biology, 1205 Docteur Penfield Avenue, Montreal, Québec, Canada, H3A 1B1; McGill University, Department Of Natural Resource Sciences, 21 111 Lakeshore Road, Sainte-Anne de Bellevue, Québec, Canada, H9X 3V9; Quebec Centre for Biodiversity Science (QCBS), McGill University, Department of Biology, 1205 Docteur Penfield Avenue, Montreal, Québec, Canada, H3A 1B1; Université du Québec à Rimouski, Département de biologie, 300 Allée des Ursulines, Rimouski, Québec, Canada, G5L 3A1

**Keywords:** Catch per unit of effort (CPUE), Drawdown, Fisheries, Flood control, Hydroelectricity, Juveniles, Littoral zone, Meta-analysis, Spawning, Trophic interactions

## Abstract

River impoundments are commonly cited as key disturbances to freshwater aquatic ecosystems. Dams alter natural hydrological regimes, homogenize river system dynamics at a global scale, can act as barriers for migratory species and may facilitate species invasions. In this synthesis, we examined the short- and long-term effects of impoundment on fish population dynamics and community structure. At the population level, we tested the “trophic surge hypothesis”, which predicts a hump-shaped response of fish abundance through time after impoundment. We tested the hypothesis on 40 recruitment time series and 125 adult abundance time series from 19 species and nine reservoirs distributed in temperate and boreal regions. At the community level, we compared diversity metrics (richness, evenness, diversity) on two datasets: 1) between reservoirs and reference ecosystems (lakes, rivers, and streams) and 2) over time (before and after impoundment and over time). At the population level, the trophic surge hypothesis was supported in more than 55% of the time series but we observed significant variation across species, reservoirs and regions. Fish recruitment increased substantially during reservoir filling and shortly after impoundment, and was usually followed by an increase in adult fish. The surge was transient and vanished after 3-4 years for recruits and after 10 years for adults. However, we are lacking long time series to conclude about population patterns in the trophic equilibrium phase. At the community level, we did not find any strong directional patterns in species diversity metrics when comparing reservoirs to reference lakes but found higher diversity and evenness in reservoirs and impounded streams/rivers relative to unimpounded streams/rivers. We did not find directional patterns when looking at a change over time. Variability in the reported diversity results across studies may be related to the ability to tease apart the unique effects of impoundment and water regulation from other stressors such as propagule pressure and eutrophication, as well as the comparability of the reference system. In conclusion, fish populations benefited quickly but transiently from impoundment, and longer time series are needed to conclude about population dynamics and equilibrium in aging reservoirs in order to develop management recommendations.

## Introduction

The creation of impoundments, and subsequent modifications of natural hydrological regimes by dams, is commonly cited as a key disturbance to freshwater aquatic ecosystems and their biodiversity (Nilsson *et al*. 2005; Dudgeon *et al*. 2006; Poff *et al*. 2007; Vörösmarty *et al*. 2010). The number and size of dams have increased considerably in the last 60 years. Construction of dams peaked in the 1960s and 1970s in boreal and temperate regions (Rosenberg *et al*. 2000) but the rate of construction is still increasing in emerging economies (*e.g*., China, Central and South America; Zhong and Power 1996; Anderson *et al*. 2006; Stickler *et al*. 2013; Li *et al*. 2013). Recent estimates suggest that 85% of the largest river systems in the United States, Canada and Europe are altered by dams (Dynesius and Nilsson 1994; Nilsson *et al*. 2005) for hydroelectricity production, irrigation, supply of freshwater, spring floods control, and recreation (Pringle *et al*. 2000; Anderson *et al*. 2006).

By transforming large rivers into storage reservoirs and by modifying lakes’ morphology (Rosenberg *et al*. 2000; Renöfalt *et al*. 2010), dams alter the natural hydrological regime of aquatic ecosystems and homogenize streamflow dynamics at a global scale (Poff *et al*. 2007). At a more local scale, upstream of dams, the regulation of the hydrological regime creates new lentic habitats (Friedl and Wüest 2002; Haxton and Findlay 2009), and generate variation in water levels far beyond natural amplitudes by imposing significant winter drawdown and by dampening spring flooding (Kroger 1973; Zohary and Ostrovsky 2011). Downstream of dams, seasonal and interannual streamflow magnitude and variability are generally reduced (Friedman *et al*. 1998; Graf 2006). These hydrological modifications, and the ways in which reservoirs are managed, can change the water temperature regime and ice cover dynamics, can impact sedimentation processes and nutrient cycling (Baxter 1977; Ostrofsky 1978; Benson 1980), can expose the littoral zone to desiccation, freezing and erosion from water drawdowns (Gaboury and Patalas 1984; Furey *et al*. 2006), and can change the general riverscape connectivity by fragmenting the fluvial network. These alterations will consequently favor the persistence of certain species over others (Stanford *et al*. 1996; Zohary and Ostrovsky 2011) by modifying the quality, diversity, and distribution of habitats, by impeding the movement of migratory species (Raymond 1979; Jansson *et al*. 2000; Morita and Yamamoto 2002; Lundqvist *et al*. 2008), and by increasing the susceptibility to non-native species colonization (Gehrke *et al*. 2002; Johnson *et al*. 2008). As yet, there has been no quantitative synthesis or general effort to understand the short- and long-term effects of impoundment on fish population dynamics Likewise, this paper examines the generality of fish α-diversity responses in reservoirs at a geographical scale much larger than previously considered.

To understand and draw generality about the effects of impoundment on fish population dynamics, scientists proposed the “trophic surge hypothesis”. In its early development, this hypothesis described and made predictions about how the increase in phosphorus loading in response to leaching and decomposition of organic matter from inundated terrestrial vegetation and soils would affect productivity in reservoirs over time (Baranov 1966; Ostrofsky 1978; Ostrofsky and Duthie 1980; Grimard and Jones 1982; Straškraba *et al*. 1993). The hypothesis has been slightly modified to predict the effects of impoundment on the overall ecosystem and on fish (Baxter 1977; Kimmel and Groeger 1986). During reservoir filling and shortly after impoundment, the increase in available littoral habitat and productivity (bacteria and algae from the increase in phosphorus) should be follow by an increase in zooplankton and macroinvertebrates and then by fishes (recruits first, followed by adults, the non-equilibrium trophic surge; Figure 1). This trophic surge is suggested to be transient and its duration will vary depending on the focal taxon’s life history traits, the degree of alteration in the hydrological regime, the amount of nutrient loading, and reservoir characteristics (Kimmel and Groeger 1986; Straskraba *et al*. 1993). The surge is followed by a non-equilibrium trophic depression in which the expansion of suitable habitats stops or decreases due to increased sedimentation and shoreline modification (Benson 1980), and where food and prey decrease (Dettmers and Stein 1992; Garvey and Stein 1998). The depression should be followed by a new trophic equilibrium where fish populations stabilize at values lower than, comparable to, or higher than before impoundment. The time to reach this new equilibrium and its state compared to before impoundment will likely depend on the focal taxon’s life history traits, the habitat suitability of the newly created reservoir, the strength of trophic interactions among species, and on other external factors such as stocking, species introductions, fishing, and mitigation measures.

**Figure 1.**
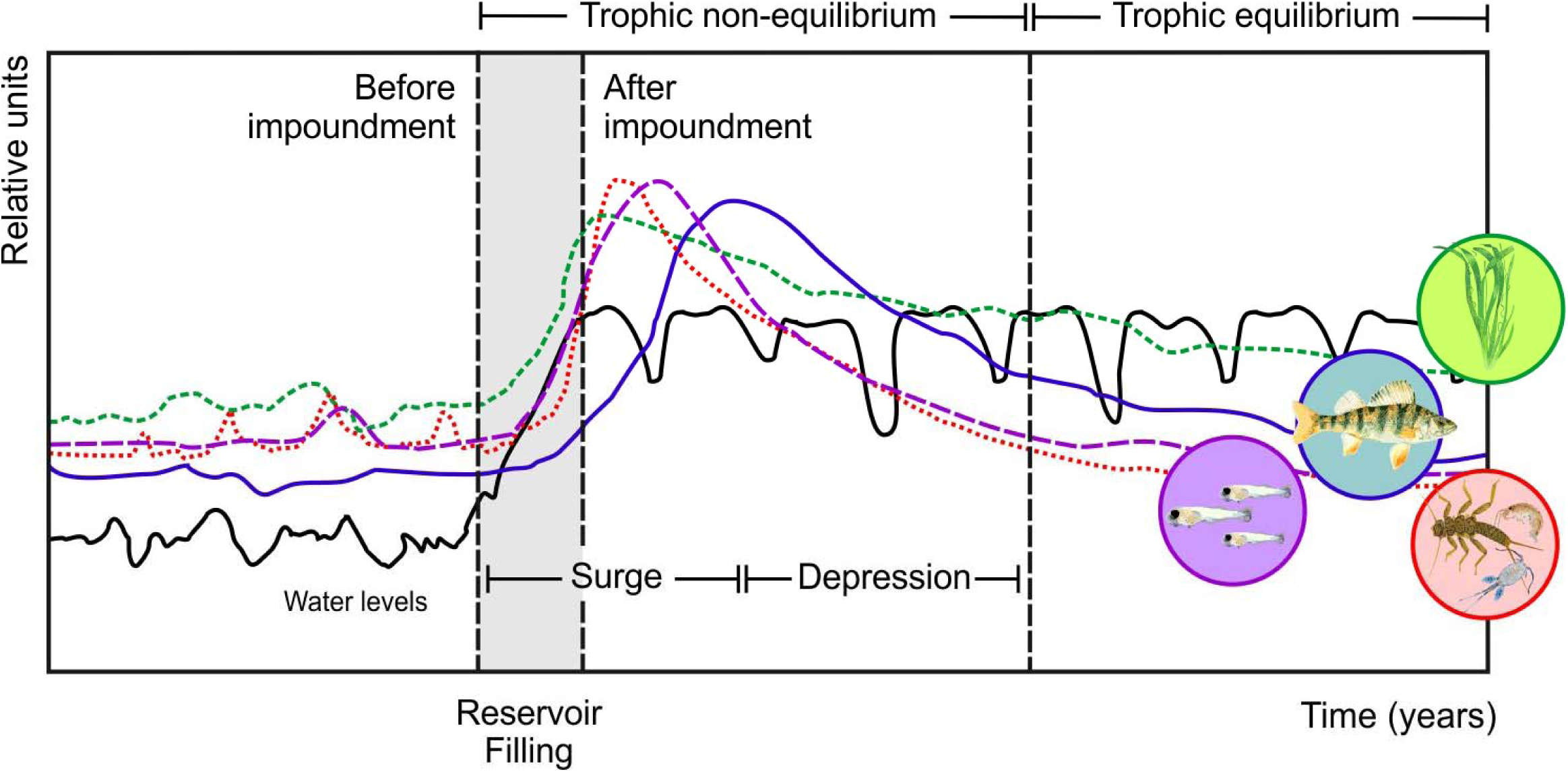
A schematic representation of the Trophic Surge Hypothesis, inspired by Kimmel & Groeger (1986). Three phases should be observed during reservoir filling and after impoundment. The first phase is a non-equilibrium trophic surge (*i.e*. increase in general ecosystem productivity), the second phase is a non-equilibrium trophic depression (*i.e*. decrease in productivity) and a third phase is a trophic equilibrium where general ecosystem productivity can stabilize to levels similar to, higher to or lower to pre-impoundment levels. On the schema, water level is represented by the black line. The available habitat for aquatic organisms (green dashed line), the abundance of zooplankton and macroinvertebrates (red dotted line), fish recruitment (purple dashed line) and adult fish abundance (blue line) are represented with relative units for figure simplicity.

Changes in quality, diversity, and distribution of newly created habitats after impoundment in reservoirs should be accompanied by changes in fish communities as well. As a general pattern, alterations of the hydrological regime and the global homogenization of streamflow by dams (Poff *et al*. 2007) are suggested to reduce the γ-diversity by causing a biotic homogenization or a convergence of fish fauna in reservoirs and impounded systems (Rahel 2000; Gido *et al*. 2009; Vörösmarty *et al*. 2010; Clavero and Hermoso 2010; Liermann *et al*. 2012). At a more local scale (*i.e*. α- and β-diversity), reservoirs and impounded systems have been reported to experience a gradual change in the fish assemblage toward lentic, generalist, invasive and non-native species (Friedl and Wüest 2002; Johnson *et al*. 2008; Haxton and Findlay 2009; Clavero and Hermoso 2010; Zohary and Ostrovsky 2011). In addition, large or untimely water level fluctuations in reservoirs are suggested to be especially detrimental for littoral and benthivorous fish species but may benefit piscivorous species by concentrating prey and predator together (Hulsey 1956; Ploskey 1986; Nordhaus 1989; Sutela and Vehanen 2008).

These initial ideas about reservoir dynamics and transient responses of fish to impoundment have been based on general observations from studies often looking at a single reservoir or a small number of comparative reservoirs over time or from more sites but over very few years. The overarching goal of this quantitative synthesis is to understand the short- and long-term effects of impoundment on fish population dynamics and community. To do so, we 1) tested the trophic surge hypothesis by examining temporal trends in fish abundance from pre-impoundment, through reservoir filling, to post-impoundment for recruitment and adult fish across species, reservoirs and sites and 2) determined how fish α-diversity, measured by the richness, diversity and evenness, and fish assemblage differ between reservoirs and reference sites (*i.e*. natural adjacent lakes and rivers/streams) and 3) determined if α-diversity change through time in reservoirs.

## Methods

### Literature search process

The studies presented in this synthesis were compiled from journals indexed in Thomson ISI’s Web of Knowledge (mostly peer-reviewed articles) and from Google Scholar (*i.e*. peer-reviewed articles and textbooks, as well as government and industry reports, non-peer reviewed journals and conference proceedings; which will be hereafter called the grey literature). Extensive searches were performed between October 2014 and June 2015 on the references available at that time. We searched for references including the following keywords, individually or in combination: “water level*”, “drawdown”, “impoundment”, “water fluctuation*”, “water regulation”, “hydrological regime” and “reservoir*” but the search included “fish*” at all times. The “*”is a meta-character and represents one or more other characters after the search terms (*e.g*., fish* can be used for fish, fishes or fishery). In addition, the reference lists and bibliographies of relevant sources were also examined to find literature that was not identified through Thomson ISI’s Web of Knowledge and Google Scholar. We then screened our database to refine our selection criteria and we included only reservoirs from temperate and boreal zones that had quantitative evidence for the effects of impoundment and water level fluctuations in reservoirs (data from figures, tables or databases in appendices) on fish population dynamics and community structure. We excluded modelling and simulation exercises from this quantitative synthesis.

### Data extraction

A total of 82 studies met our search criteria. For each study, we consistently recorded: 1) the temporal scale examined (*i.e*. whether the data came from before, during and/or after impoundment, as well as whether the study examined the impacts associated with inter- and intra-annual variation in water levels), 2) whether the study was conducted at the population or fish community level, 3) whether the study was from the peer-reviewed or the grey literature, 4) the number of replicates (*i.e*. the number of reservoirs being compared) and 5) the number of reference sites (*i.e*. adjacent natural lakes/rivers/streams).

From the 82 studies, we obtained data on a diverse set of reservoirs (n = 154) but site-specific information was not provided in two studies that had done an extensive comparison (n > 100 across both studies) of fish communities in reservoirs and reference ecosystems (*i.e*. lakes, rivers and streams) (Irz *et al*. 2006; Gido *et al*. 2009). In all other cases, we were able to extract the following set of information from each study: 1) the coordinates of the reservoirs and of the references lakes, rivers or streams when available, 2) the area of the reservoir and reference lakes (ha), 3) key dates including the date at which the reservoir was created, the date at which the river or lake was impounded and the date at which the reservoir reached its full pool, and 4) the number of years of data in the time series.

From our initial dataset of 82 studies, we used a subset of 30 that met our analysis criteria to evaluate the effects of impoundment on fish population dynamics and community structure. Studies with only a few years of data at the population level (recruitment or adult abundance) or lacking information before impoundment cannot be used to test the trophic surge hypothesis, and therefore were excluded from our quantitative synthesis. From these 30 studies, seven had suitable time series covering the period before and after impoundment to test the trophic surge hypothesis (Tables A1 and A2 in Appendix A). Within these seven studies, we identified and extracted 40 continuous time series examining fish recruitment for 18 species, and 125 time series examining adult fish abundance for 19 species. These time series were collected from eight different reservoirs (19 sites; some reservoirs had more than one sampling sites) and lasted between eight and 23 years long (Tables A1 and A2). Most of the time series, especially the longer ones, were collected for species of recreational fisheries interest (*e.g*., northern pike, yellow perch, walleye and sauger). From the 30 studies, we also extracted data from 23 studies to evaluate the effect of impoundment on the fish community. In developing the database on fish communities, we classified the studies based on their experimental design. In some studies, data were available comparing reference natural systems and reservoirs communities (*i.e*. snapshot in time: few years and many sites) or looking at before and after impoundment (*i.e*. snapshot in space: many years and few sites).

Data about abundance or relative abundance for a given species were mostly extracted from tables presented in the articles and reports. When data were only presented in figures, they were extracted using the GetData Graph digitizer software (v. 2.26). Some data were also extracted from supplemental material available online.

### Data analysis

#### Testing the trophic surge hypothesis for recruitment and adult abundance

*General trend in the time series* – To test if the trophic surge hypothesis was supported by the available datasets, we compared the fit of eight alternative functions corresponding to general abundance patterns that could be observed for the period covering before impoundment, during reservoir filling and after impoundment (Table B1 in Appendix B). We tested two hump-shaped patterns suggested to represent the trophic surge hypothesis (Ricker and negative quadratic polynomial functions), one “U”-shaped pattern where abundance is high before impoundment, decrease during filling or shortly after and then come back to high abundance values (positive quadratic polynomial function), two non-linear decreasing patterns (negative exponential and quadratic polynomial functions), one non-linear increasing pattern (exponential function), one linear increasing pattern, one linear decreasing pattern and a null or no pattern (flat line function). Parameters used to fit each function to the observed data were estimated by maximizing the normal Gaussian likelihood using the mle2 function with the Nelder-Mead method available on the bbmle package version 1.0.4.1 (Bolker and R Development Core Team 2012). To evaluate which function fit best the observed data, we used Akaike’s Information Criterion modified for small sample sizes (AICc; Tables B2 and B3). We calculated the difference between AICc for each model *i* and the lowest observed AICc (∆AICc), and compiled normalized Akaike weights (*w_i_*) for each function to estimate the probability that function *i* is the best function, given the data and set of candidate functions (Burnham et al 2011). All analyses were performed in R (*v*. 3.2.0). If the trend was hump-shaped, the value of the peak was extracted by dividing the actual value at the peak for a given species in a given reservoir by the mean value of abundance before impoundment. We also extracted the time at which the peak occurred in time in relation to time since impoundment (0 = dam is operational and the reservoir reached full capacity). We used two criteria to determine if the equilibrium phase was reached for a given time series. First, the time series should not demonstrate an increasing or a decreasing trend for at least four consecutive years at the end of the time series and second, the coefficient of variation (CV) of the abundance in the last four years of the time series should be comparable to the CV of the abundance before impoundment (*i.e*. within 1 order of magnitude).

#### Calculation of the diversity metrics

*Richness –* In 20 of the 23 studies identified, we were able to estimate total richness (total number of species sampled) in reservoirs and reference ecosystems from the available data about fish relative abundance. In the other studies, only rarefied richness estimates were available (Głowacki and Penczak 2000; Gido *et al*. 2009) or total richness with no information about fish relative or total abundance (Edwards 1978). *Diversity* – From the 20 studies that provided relative fish abundance or total abundance, we were able to calculate diversity by using the Shannon-Weaver index. *Evenness* – From the 18 studies that provided relative fish abundance or total abundance, we were able to calculate evenness by using the Pielou index. *Percentage of littoral and benthivorous fish in the community* – The impacts of water-level regulation, particularly winter drawdown in reservoirs are often described as being most evident in the littoral zone. This suggests that littoral and benthivorous fish species could be especially vulnerable because access to their preferred habitat becomes limited and zoobenthic food sources can be reduced (Furey *et al*. 2006; Sutela and Vehanen 2008). We first categorized fish species based on their feeding and spawning habitat, extracting information provided by FishBase (Froese and Pauly 2015), Scott and Crossman (1973) and categorizations already made by some studies (Gendron 1990; Sutela and Vehanen 2008). We then calculated the proportion of littoral and benthivorous fish species in each reference lake and reservoir. *Species turnover rate* - Jaccard’s dissimilarity distance metric was applied to quantify the degree of species turnover after impoundment (Anderson *et al*. 2011). To extract the dissimilarity matrix from each reservoir, we used the vare.dist function with the Jaccard distance in R (vegan library v. 2.3-0). Because the focus of our analyses were to define the difference in diversity before and after impoundment, we used only the first column of the dissimilarity matrix.

#### Effects of impoundment on fish community and assemblages

We asked four questions regarding the impact of impoundment on the structure and diversity of fish communities based on the available datasets. First, we asked whether community structure metrics differed from references sites (lakes in seven studies and streams in two studies). For this question, we also examined if fish assemblage, measured by the percentage of littoral and benthivorous fish, differed between reservoirs and reference lakes. For the comparison of reservoirs and reference ecosystems we used mini random effects meta-analyses. For richness, we performed three mini meta-analyses using 1) references lakes and references rivers/streams together, 2) reservoirs and references lakes and 3) reservoirs and impounded streams and reference streams and rivers. For diversity and evenness, we performed a mini meta-analysis comparing fish community in reservoirs and reference lakes because data were not available from the two studies comparing reference rivers/streams and impounded streams and reservoirs. Mini meta-analyses were performed using the package metafor (v. 1.9-7; Viechtbauer 2010). Effect sizes were calculated as Hedges’ *d* (Hedges 1981; Hedges and Olkin 1985) to estimate standardized difference between the mean values in reference ecosystems and reservoirs, according to their standard deviations and sample sizes. A positive effect size for a given metric indicates higher values in reservoirs relative to reference ecosystems. Heterogeneity of effect sizes was examined with *Q* statistics (Hedges and Olkin 1985), which can be used to determine whether the variance among effect sizes is greater than expected by chance. For the comparison of diversity and evenness between reservoirs and reference streams/rivers, we extracted mean diversity and mean evenness in reservoirs and streams from three river basins (supplemental tables S1 and S2 in Gido *et al*. (2009); Missouri basin, 9 reservoirs:54 streams; Arkansas, 14:73 Red, 5:16). We than ran General Linear Mixed effects Models (LMM) with the restricted maximum likelihood method (lmer function from the lme4 library, v. 1.1-7 in R). We evaluated support with 95% CI. As a second question, we asked whether richness, diversity and evenness changed before vs. after impoundment. If fish communities were sampled over multiple years, we first averaged richness, diversity and evenness values before and after impoundment (two categories) and then compared them using a LMM. To account for variation and specificity related to each study, we nested individual reservoirs (L_R variable) within each study as random factors (L_R|Study ID). All variables were z-standardized prior to analysis. Third, we asked how richness, diversity and evenness changed over time (*i.e*. increase, decrease or no change) by using all the data points spanning before, during the filling phase and after impoundment. For this question, we also used a LMM with study ID used as a random factor. Finally, we asked whether species turnover rate (Jaccard’s index) changed through time following impoundment using LMM (Study ID was used as a random factor).

## Results

### Summary of the datasets

From the 82 studies that met our search criteria, 43% of the 154 reservoirs were located in Canada, 41% in the United States and 16% in Europe (Figure 2). We found that the reservoirs studied varied widely in age, area, and in temporal and spatial extent of associated data. The vast majority of reservoirs were created between 1930 and 1980 with a peak around 1960 (Figure 3a). They varied extensively in surface area ranging from 50 to 427 500 ha (Figure 3b). On average, field data collected to examine the effect of impoundment and water regulation on fish were collected over 9 years (Figure 3c). However, the frequency distribution of the length of the time series is clearly bimodal, highlighting that for many studies, just a single year’s worth of data was collected. These single year of data reservoirs were often part of a larger comparative study (*i.e*. comparing fish communities among numerous reservoirs and reference lakes and rivers). A substantial proportion of the studies (36%) were published in the grey literature.

**Figure 2.**
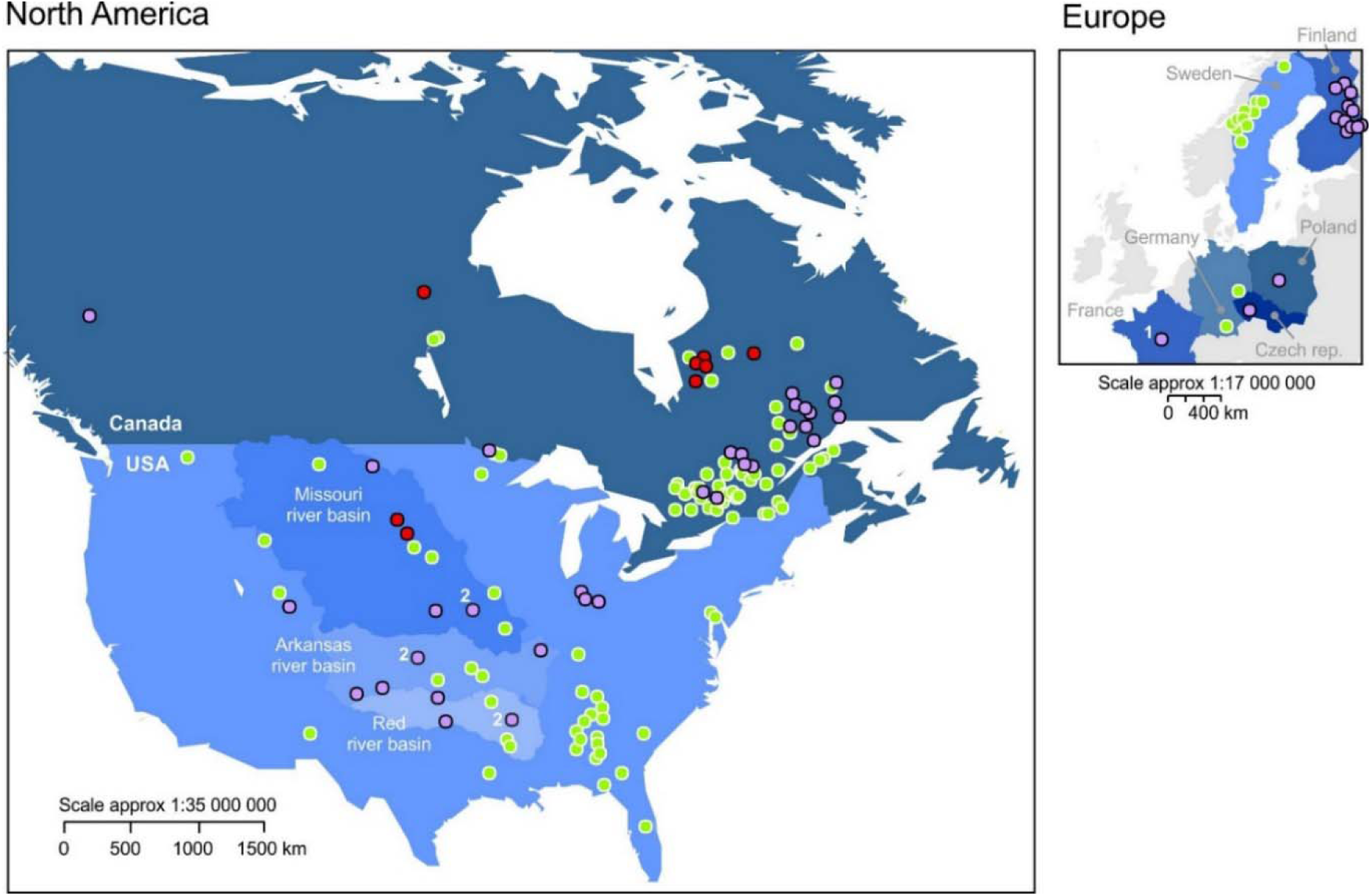
Spatial distribution of the 154 reservoirs from 82 studies examining the effects of impoundment and water level variation on fish in boreal and temperate regions of the Northern Hemisphere. Red dots represent the reservoirs on which we extracted quantitative data to test the Trophic Surge Hypothesis and the purple dots represent the reservoirs on which we extracted quantitative data to evaluate the effects of impoundment on fish richness, evenness and diversity. Some of the sites on the maps refer to studies where multiple sampling locations (> 100) were covered in a single study, without detailed information provided in the original study about specific site characteristics (Irz et al. 2006 and Gido et al. 2009). These studies on the map have superscript numbers.

**Figure 3.**
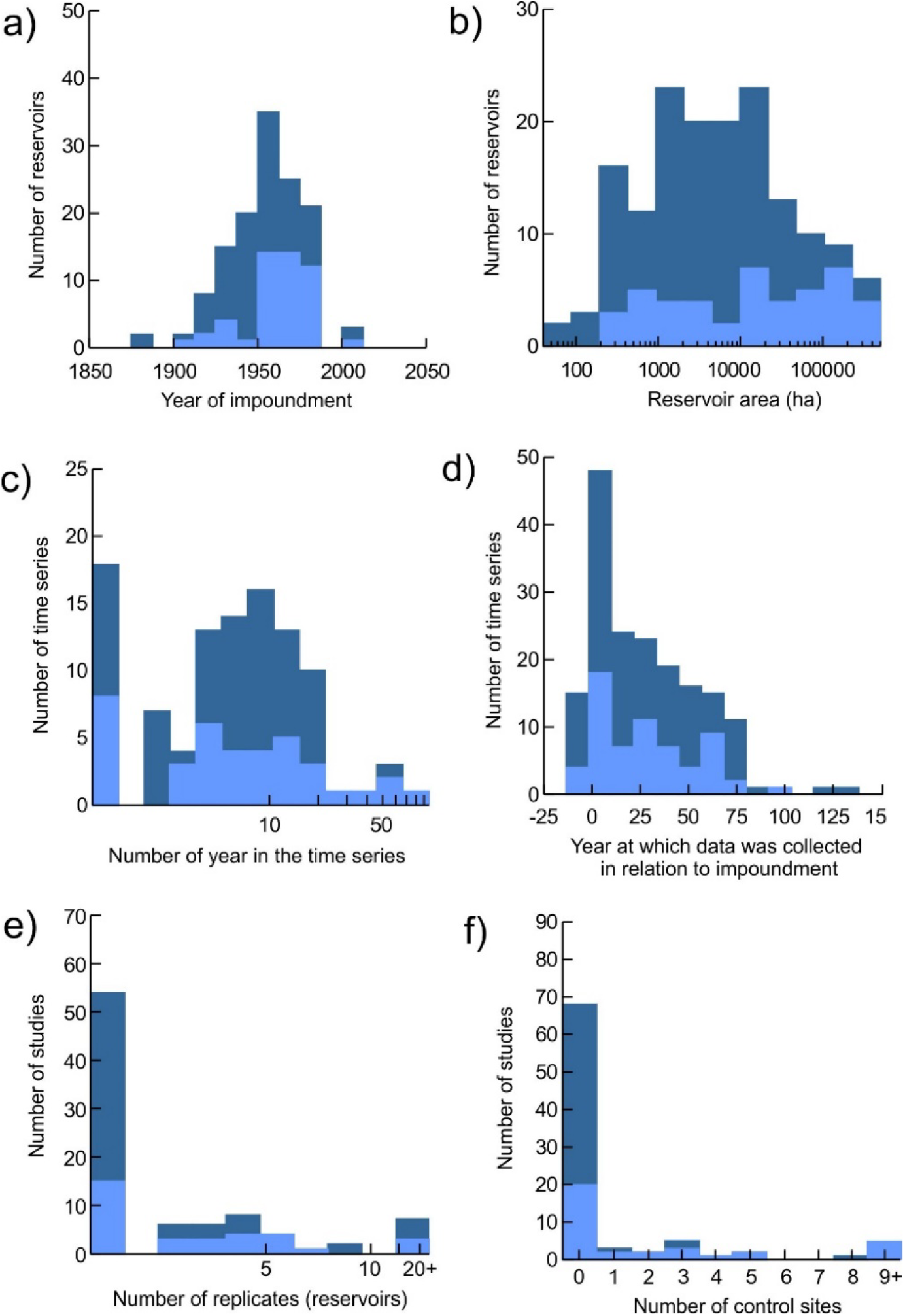
Frequency distributions of the number of reservoirs studied in relation to a) year in which the river or lake was impounded, b) the size of the reservoirs (ha). Frequency distribution of the number of time series in relation to c) the number of years in the time series and d) the year at which data was collected in relation to impoundment (minus values represents time series in which we have data before impoundment). Frequency distribution of the number of studies in relation to e) the number of replicates (reservoirs) and f) the number of control sites (un-impounded streams, rivers and lakes). The dark blue bars represent the overall dataset meeting our search criteria and the light blue bars represent the dataset used in this quantitative synthesis to test the trophic surge hypothesis and to evaluate a change in community structure following impoundment.

We identified a subset of studies (N=30) that met our analysis criteria and thus enabled us to test the trophic surge hypothesis and to evaluate how impoundment affects fish community structure. This subset of studies also had a broad geographic distribution, as 39% of the 44 reservoirs were located in Canada, 37% in the United States and 23% in Europe (Figure 2; black outlined dots). We found that this subset of studies (light blue bars; N=30 studies, 44 reservoirs) had very similar frequency distributions to the full datasets (Figure 3; dark blue bars; N=82 studies, 154 reservoirs), and thus could be considered as a representative subsample. We identified 13 studies in which time series were collected to cover the periods before and after impoundment (bars on the left side of the green dashed line; Figure 3d). However, for most studies, field observations began many years after impoundment (Full dataset, median = 22 years, Subset, median = 18 years). As a general pattern, very few studies examined the effects of impoundment across several reservoirs (number of replicates; Full dataset and Subset, median = 1, range= 1 to 23; Figure 3e) and even fewer had reference sites (adjacent un-impounded lakes, rivers or streams; Full dataset and Subset, median = 0; range = 0 to 8, Figure 3f).

### Is the trophic surge hypothesis supported by the data?

#### Recruitment of young of the year

More than fifty percent of the time series (55%; 22/40 time series) showed a clear increase in fish recruitment or very high recruitment values during and shortly after reservoir filling (*i.e*. the surge) relative to the pre-impoundment period, and/or showed a decrease in recruitment after impoundment (*i.e*. the depression) which support the trophic surge hypothesis (Table A1 and Table B2). From the 40 time series, 43% showed a clear hump-shaped pattern (17/40 time series; Figure 4a), 20% showed an increasing trend (8/40 time series; Figure 4a), and 15% showed a decreasing trend (6/40 time series; Figure 4a). From the time series showing an increasing trend, only one was in response to impoundment (not enough data points after impoundment to capture statistically the depression phase) and the others cannot be attributed to impoundment (7/40, Figure 4a, Table A1). From the time series showing a decreasing trend, four actually captured the end of the trophic surge and the depression phase and were in response to impoundment but two cannot be clearly attributed to impoundment (Figure 4a, Table A1) We were unable to statistically detect a pattern in 22% of the time series (No pattern; 9/40, Table A1 and Table B2). From the 32 time series on which we performed the curve fitting analysis, 50% showed strong support for hump-shaped patterns based on AICc scores (16/32 time series; Tables A1 and B2). The Ricker hump-shaped pattern had a higher Akaike weight than the polynomial hump-shaped pattern (*w_i_*; 0.295 vs. 0.215; Table B2) suggesting that the peak in recruitment occurred early in time.

**Figure 4.**
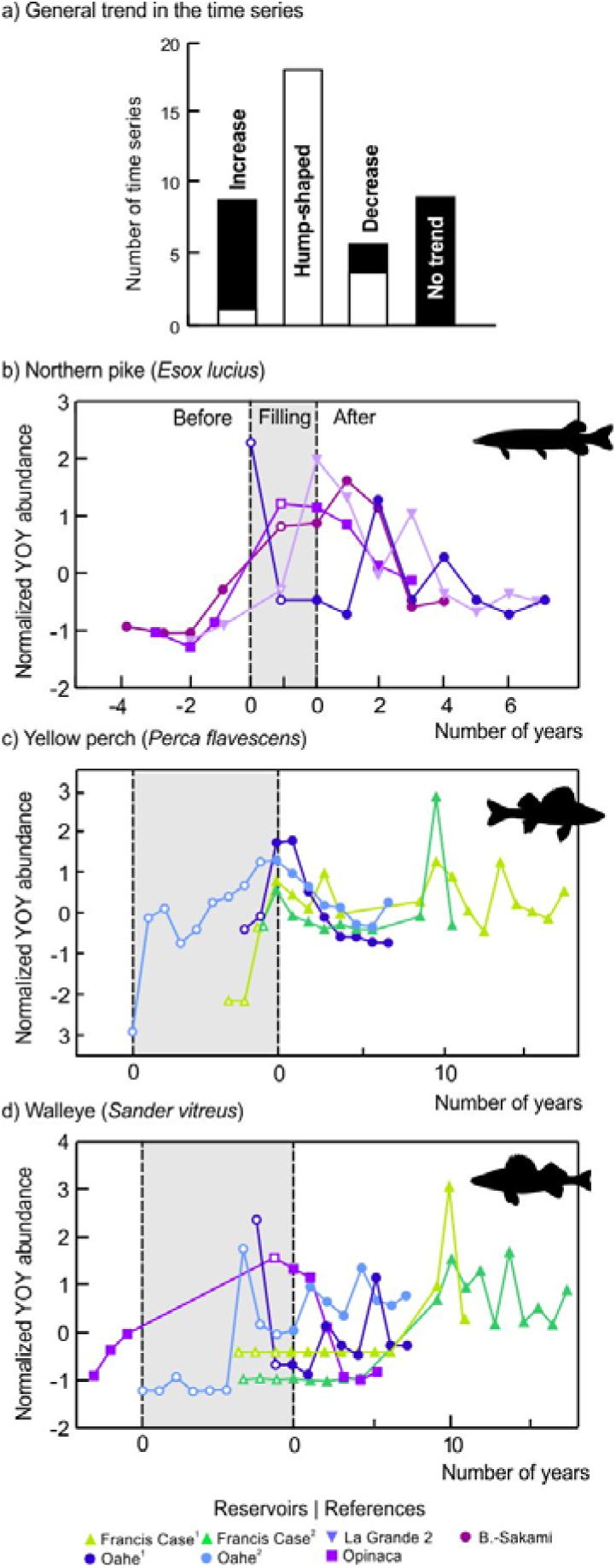
a) Summary of the number of species-by-reservoir time series for which recruitment (young of year fish abundance; YOY) showed increasing, hump-shaped, decreasing, or no trend across the period of record. The white shaded area in the increase and decrease bars represents the number of time series supporting the trophic surge hypothesis graphically, but for which we had limited data before and/or after impoundment to statistically detect a hump-shaped pattern with the curve fitting analysis. The black shaded area represents the number of time series that do not support the trophic surge hypothesis, graphically and statistically. b), c), and d) are example time series of recruitment for northern pike, yellow perch, and walleye before impoundment, during filling (represented by the grey area with open symbols), and after impoundment

When a hump-shaped pattern was detected, recruitment was 4.3 ± 3.2 (mean ± SD) times higher during filling and shortly after impoundment relative to before impoundment (magnitude of the peak; range = 1.2 to 17.1 times higher; Table A1). The peak occurred 0.4 ± 1.9 years after the dam was operational (Table A1). The effect of impoundment was transient, vanishing 3-4 years after impoundment for most of the hump-shaped time series (see examples in Figure 4 b, c and d). For the time series that extended over periods greater than 5 years after impoundment, few showed a clear directional change (*i.e*. higher or lower abundance compared to before impoundment). From the 40 time series, 30% showed a trend toward a higher abundance after impoundment relative to before, 25% appeared to show a decreasing trend but the difference was significant in only 9 time series (Table A1). Forty-five percent of the time series showed no trend or got back to values comparable to before impoundment. Only 25% of the time series clearly reached an equilibrium at the end of the dataset (Table A1).

We observed significant variability in recruitment patterns among species and reservoirs. For example, the northern pike showed a clear hump-shaped pattern in boreal reservoirs (Québec) but a decreasing trend in a temperate reservoir (Oahe; South Dakota; Figure 4b). It is important to note though that we do not have any observations pre-dating the creation of the Oahe reservoir because filling the reservoir took 9 years. In the Opinaca reservoir (6 months to fill), La Grande 2 (1y), and the Boyd-Sakami diversion (1y), a general increase in recruitment was observed the year at which reservoirs reached their operational level for many species (Figure C2). Thus part of the variability across time series may appear because of variation in baseline conditions (*e.g*. time to fill the reservoir).

#### Adult fish abundance

Nearly 60% of the adult abundance time series supported the trophic surge hypothesis (65/109 time series; Table A2 and Table B3). From the 109 time series, 45% showed a clear humped-shaped pattern where the abundance of adult fish increased during reservoir filling or shortly after impoundment and then decreased in the depression phase (49/109; Figure 5a), 7% showed a significant increasing trend (8/109 time series; Figure 5a), and 23% showed a significant decreasing trend (25/109 time series; Figure 5a). From the time series showing an increasing trend, three were in response to impoundment (*e.g*., showed a peak graphically shortly after impoundment but the hump-shaped curve was not supported statistically), one was not and four visually showed a “U-shaped” pattern not supported statistically (Figure 5a, Table A2). From the time series showing a decreasing trend, six were in response to impoundment, 16 were not and three visually showed a “U-shaped” pattern not supported statistically (Figure 5a, Table A2). We were unable to statistically detect a pattern in 25% of the time series (No pattern; 27/109, Table A2 and Table B3). Overall, the Ricker hump-shaped pattern had a slightly higher Akaike weight (*w_i_*; 0.249) than the polynomial hump-shaped pattern (*w_i_*; 0.225; Table B3) suggesting that the peak in abundance occurred early in the time series. Based on visual assessment, many time series for walleye and whitefish showed a “U” shaped pattern, which although not detected statistically (13/109; Table A2), depicted a trend where abundances were relatively high before impoundment, then reached low values shortly after impoundment and then increased to reach their highest values roughly 10 years after impoundment.

**Figure 5.**
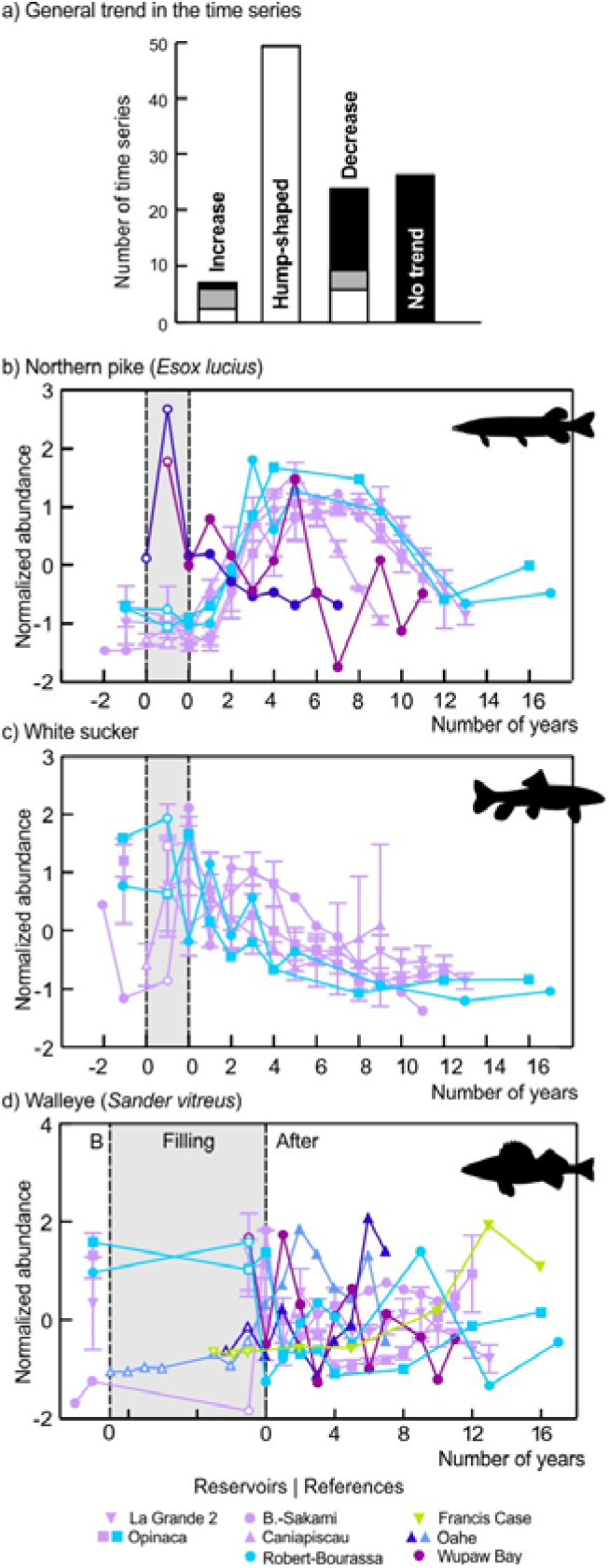
a) Summary of the number of species-by-reservoir time series for which adult fish abundance showed increasing, hump-shaped, decreasing, or no trend across the period of record. The white shaded area in the increase and decrease bars represents the number of time series supporting the trophic surge hypothesis graphically, but for which we had limited data before and/or after impoundment to statistically detect a hump-shaped pattern. The black shaded area represents the number of time series that do not support the trophic surge hypothesis, graphically and statistically. The grey shaded area represents time series that graphically showed a “U-shaped” pattern that was not detected statistically with the curve fitting analysis. b), c), and d) are example time series of normalized adult fish abundance for northern pike, white sucker, and walleye before impoundment, during filling (represented by the grey area with open symbols), and after impoundment.

When a hump-shaped pattern was detected (49/65) or strongly suspected based on visual assessment (16/65), adult abundance was 14.0 ± 22.5 times higher during filling and shortly after impoundment relative to before impoundment (magnitude of the peak; range = 1.0 to 188.6 times higher; Table A2). The peak in adult abundance occurred in average 2.3 ± 2.2 years after the dam was operational, which is lagging two years after the peak in recruitment (Table A2). The effect of impoundment on adult fish was transient, but lasted longer than for recruitment. The effect of impoundment (surge and depression) seemed to vanish 8.9 ± 4.1 years after the dam was operational but many time series were still in the depression phase (*i.e*. did not appeared to have reach the equilibrium phase) at the end of the series. Twenty-eight percent of the time series seemed to have reached an equilibrium phase at the end of the time series, including species that reached very low values and appeared to have crashed, so directional trends have to be interpreted with caution (Table A2). Overall, 25% of the time series showed significant directional change (*i.e*. higher or lower abundance after impoundment compared to before impoundment; Table A2).

Even after taking into account the differences in time series length across studies, we observed a striking consistency in trends for some taxa. As an example, northern pike showed a clear and consistent hump-shaped pattern in Québec reservoirs, peaking roughly six years after impoundment (5.5 ± 2.0 years) and then decreasing to abundance levels comparable to before impoundment after roughly 11 years (Figure 5b, Table A2). In contrast, other taxa demonstrated directional responses, where some species clearly benefited from impoundment (*e.g*. whitefish; Table A2) and others, such as the white sucker showed a consistent decline over time (Figure 5c, Table A2). Finally, walleye adult abundance showed no strong patterns over time and a lot of variation across reservoirs (Figure 5d).

### How does impoundment affect fish α- and β-diversity?

#### How do fish communities in reservoirs compare to those in reference ecosystems?

Overall, we found that species richness, diversity and evenness were comparable between reference lakes and reservoirs (Figure 6). Species richness was comparable between rivers/streams and reservoirs (Figure 6). Fish diversity and evenness were consistently and clearly higher in reservoirs relative to reference streams/rivers. More specifically, when examining studies comparing reservoirs and reference lakes from the mini meta-analyses, we observed approximately as much variation in richness and diversity within any focal study as between studies (richness = *Q* = 37.70, p< 0.001, df = 10; diversity =*Q* = 11.27, p< 0.001, df = 5), but not for evenness (*Q* = 4.597, p = 0.467, df = 5; Figure 6). From the meta-analyses comparing richness in reservoirs and reference ecosystems, we found no overall difference despite a trend toward higher richness in reservoirs (Figure 6a (all), p = 0.069), definitely no difference between reservoirs and reference lakes (Figure 6a (lakes), p = 0.596) and a slight, but not significant trend for higher richness in reservoirs relative to reference rivers/streams (Figure 6a (riv.), p = 0.089). Using, LMM, we found that diversity in reservoirs was 1.59 ± 0.09 times higher than in streams from three river basins (LMM; estimate ± SE = −0.87 ± 0.17, 95%CI = −1.05 to −0.71; t-value = −4.91). We also found that evenness was 1.48 ± 0.03 times higher in reservoirs than in streams from the same three river basins (LMM; estimate ± SE = −0.19 ± 0.02, 95%CI = −0.21 to −0.16; t-value = −8.54) The proportion of littoral and benthivorous fish inhabiting the littoral zone was comparable between reservoirs and reference lakes (Figure 6d; p=0.339), except for reservoirs and natural lakes from Finland where the proportion of littoral and benthivores fish species in the community was on average 53% in reservoirs compared to 91% in natural lakes (Sutela & Vehanen 2008).

**Figure 6.**
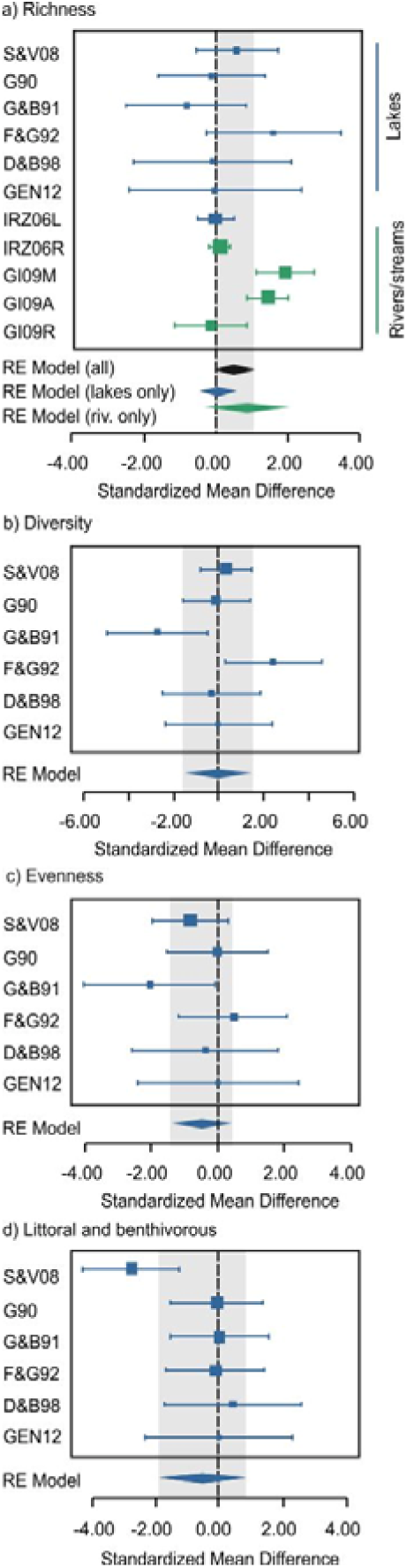
a) Meta-analysis plots for a) richness (*i.e*. the number of species), b) diversity (Shannon-Wiener index), c) Evenness (Pielou index) and e) proportion of littoral and benthivorous fish in the community. The small blue and green boxes are the standardized mean effect size (Hedge’s d) with error bars representing the bias-corrected 95% confidence intervals. Effects are significant if their confidence intervals do not overlap zero. Random effect (RE) model indicates the overall effect combining information from all studies. For richness, we evaluate the mean effect for all reference ecosystems combined (black RE model (all; lakes and streams/rivers), when reservoirs are compared with reference lakes only (blue RE model) and when reservoirs are compared with streams/rivers only (green RE model).

#### How do fish communities in reservoirs differ before and after impoundment?

Overall, there was no strong predictable pattern in richness, diversity, evenness or species turnover rate before and after impoundment or over time since impoundment with continuous data (Figure 7, Figure D1 in Appendix D). From nine studies distributed among 10 reservoirs, species richness was, on average, slightly lower after impoundment than before (Figure 7a, Table 1), but no trend was found with continuous data (*i.e*. sites were monitored every year or in a discontinuous fashion after impoundment; Figure 7b, Table 1). Diversity and evenness did not differ (Figure 7c-f, Table 1) before and after impoundment or when data were examined continuously. Species turnover rate varied widely among reservoirs, showing increasing dissimilarity over time for some reservoirs and increasing similarity for others but no significant pattern was detected overall (LMM: estimate ± SE = −0.002 ± 0.002; 95%CI = −0.006 to 0.003; Figure D1).

**Table 1.** General Linear Mixed effects models (LMM) examining the variation in richness, diversity and evenness before and after impoundment. In the first analysis, observed values were binned into “before” and “after impoundment” categories. In the second analysis, we used a finer temporal resolution when sites were monitored over 6 continuous years to 20 discontinuous years after impoundment. In all cases, we used a random effect to identify the Study ID to avoid pseudoreplication. The following statistics are reported: *t* values and the coefficient estimate with its standard error (Estimate ± SE). Models for which the 95% confidence interval (CI) did not overlap zero are indicated in bold font.

| Community metrics | t or F values | Estimate ± SE | 95% CI |
| --- | --- | --- | --- |
| *Before and after* |  |  |  |
| Richness | **1.426** | **-2.535 ± 1.778** | **-0.828 to -4.241** |
| Diversity | 0.243 | -0.037 ± 0.150 | -0.181 to 0.108 |
| Evenness | 0.212 | -0.012 ± 0.055 | -0.064 to 0.041 |
| *Finer temporal resolution* |  |  |  |
| Richness | -0.021 | -0.001 ± 0.047 | -0.039 to 0.052 |
| Diversity | 0.111 | -0.001 ± 0.004 | -0.004 to 0.003 |
| Evenness | -0.091 | 0.001 ± 0.002 | -0.001 to 0.001 |

**Figure 7.**
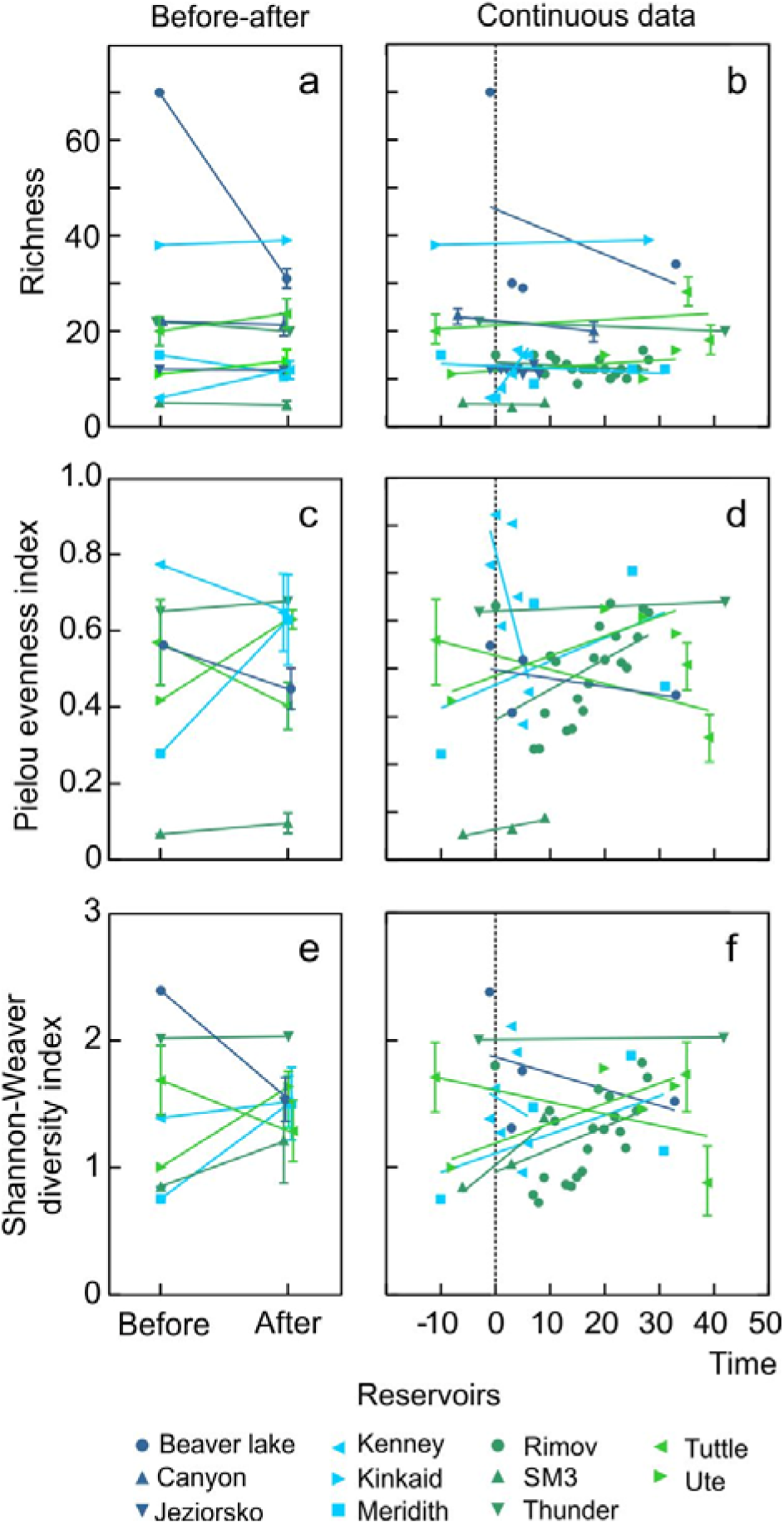
Comparison of a-b) mean richness, c-d) mean diversity (Shannon-Wiener index) and e-f) mean evenness (Pielou index) before and after impoundment and over a finer temporal resolution, across 13 boreal and temperate reservoirs. In the before and after panels (a,c,e), when multiple years of data were available either before or after impoundment, they were binned (mean effect ± SE) in order to get one value before and one value after impoundment.

## Discussion

This synthesis is the most comprehensive work that quantitatively evaluates the effects of impoundment and reservoir aging on fish population dynamics and α-diversity in the northern hemisphere. Two major insights emerge from our analysis. First, by examining nearly 170 recruitment and adult abundance time series from 9 reservoirs and 19 species, we found good support for the trophic surge hypothesis, mainly for the non-equilibrium phase in the early years following impoundment (*i.e*. trophic surge followed by a depression phase; Baranov 1966; Kimmel and Groeger 1986; Straškraba *et al*. 1993). However, the paucity of long time series after impoundment impeded strong conclusions about the state of fish populations in the equilibrium phase. Second, by examining reservoirs distributed from the temperate to the boreal ecozones, we did not find strong empirical evidence for a decrease in species α-diversity when we compared richness, diversity and evenness in reservoirs and reference lakes or when looking for a change in α-diversity over time, following impoundment (*i.e*. before and after data and continuous data). However, we did find higher diversity and evenness in reservoirs when compared to adjacent reference rivers and streams. Our findings demonstrate that monitoring fish populations and community for only a few years after impoundment, during the non-equilibrium phase, could give a false impression of a highly productive ecosystem and may lead to erroneous management recommendations. Our findings also suggest that the homogenizing effect of impoundment on fish fauna might vary from one region to the other (*e.g*., not significant or detectable in boreal ecoregions) and might be the result of several synergistic factors that are confounded with impoundment (e.g. propagule pressure).

### The trophic surge hypothesis is generally supported by our synthesis

#### Good support for the trophic non-equilibrium phase but little support for the equilibrium phase

The trophic surge hypothesis is a useful and plausible framework to predict how fish populations respond to impoundment and reservoirs aging. The hypothesis had good support to explain recruitment and adult abundance during the non-equilibrium phase (*i.e*. trophic surge and depression) during. Our results from recruitment time series demonstrated that fish reproduction and larval survival were likely very successful during reservoir filling and shortly after impoundment, which can largely be attributed to an increase in the availability of spawning and nursery habitats after impoundment. A peak in recruitment was followed by an increase in adults, lagging behind by 2-3 years. The effect of impoundment was transient and seemed to vanish after roughly 3-4 years for recruits and after roughly 8 years for adults, where abundances were back to pre-impoundment values. However, the paucity of long time series after impoundment impeded our ability to develop strong conclusions about the actual abundance values observed at equilibrium and the time needed to reach such a trophic equilibrium phase. The longest time series covered a maximum of 15 years after impoundment for recruitment and 17 years for adult abundance. For the adult fish dataset, 72% of the time series did not convincingly reach the equilibrium phase and were still increasing or decreasing at the end of the time series. Likewise, it is difficult to conclude about a change in average abundances after impoundment relative to before when we only have 2-3 data points in the period before impoundment (85% of the time series; Table A2).

Two studies with longer time series (>40 years; Cohen & Radomski 1993; Hendrickson & Power 1999) showed that the time needed to reach equilibrium after impoundment can vary strongly among species and reservoirs. These studies were not included in our synthesis because they lacked data before impoundment or presented commercial catches as a proxy for fish abundance. Nonetheless, Hendrickson and Power (1999) showed that the mean yearly total CPUE for all species combined in the Sakakawea Reservoir was very high during the first five years of filling, then declined sharply and stabilized eight years after impoundment at values three times lower than values observed during filling. They also showed that the abundance of crappies and goldeye followed a hump-shaped trajectory, stabilizing 10 years and eight years after the start of reservoir filling, respectively. The walleye, whitebass and Johnny darter demonstrated an increasing trend but they were introduced or stocked. The yellow perch and the common carp demonstrated a decreasing trend where they were at high abundance during the first years of filling and then dropped sharply 10 years and five respectively after the start of reservoir filling. Twelve other species did not show any significant trend in relation to impoundment (Hendrickson and Power 1999). In Rainy (impounded in 1909) and Namakan reservoirs (impounded in 1914), 72 years of data of lake fish commercial catch showed that the species reached a stable equilibrium quickly after impoundment. Commercial catch of walleye was very high following impoundment and decreased consistently since the 40s (Cohen and Radomski 1993). Unfortunately, CPUEs trends (not measured until 41 years after impoundment) showed very different patterns than the catch data and thus this brings into question how representative are the catch time series of fish abundance trends.

#### What are the potential sources of variation for the observed patterns?

When time series from all reservoirs and species were combined for recruitment and adult abundance data, respectively (Figure C1), or when data across reservoirs were combined for a given species (Figure C2), our analyses demonstrated the absence of a trend. This lack of support for a general pattern suggests that there is considerable heterogeneity at the level of individual studies, reservoirs and species.

We have been sensitive to the fact that length of the time series and the baseline information about reservoir history and management (*e.g*. time needed to fill the reservoir) may influence ones’ interpretation of time series and the support for the trophic surge hypothesis. Time needed to fill the reservoirs varied quite extensively among studies, going from nine years in Oahe reservoir (Nelson and Walburg 1977) to only six months in Opinaca (DesLandes *et al*. 1995). Figure 4c suggests that yellow perch recruitment in Oahe reservoir peaked after the reservoir reached its operational level and got back to pre-impoundment values after four years. However, a much longer time series from this same reservoir and covering nine years before impoundment (*i.e*. at the start of reservoir filling) and eight years after, clearly showed that yellow perch recruitment increased drastically right after the start of reservoir filling, and remained at values much higher than pre-impoundment values during filling and after impoundment (Figure 4c). A similar pattern can be observed in walleye (Figure 4d). Unfortunately, across the 44 time series, less than 30% of the time series (N = 13, Tables A1 and A2) covered the periods before filling (*i.e*. 1 or 2 data point before filling), during filling and after impoundment (*i.e*. at least 5 years of data after filling). Even with this more select group of time series, our ability to appropriately characterize the pre-impoundment conditions was limited given that fish recruitment is highly stochastic, fluctuates among years and is influenced by density-dependent and density-independent factors (Myers 1998, 2002; Karjalainen *et al*. 2000; Grenouillet *et al*. 2001). However, given that the hump-shaped pattern was predominant among the adult time series which are more stable, we feel justified in concluding that there is support for the trophic surge hypothesis overall.

The variation observed among reservoirs and species can also arise from external and internal abiotic and biotic factors, from reservoir characteristics and its management (*e.g*., mitigation of the impacts by the industry and stocking), and from fisheries activities. These site-specific effects can confound or mask the effect of impoundment. For example, DesLandes *et al*. (1995) suggested that fish movement and redistribution following impoundment can explain some fish species’ abundance patterns at a given site. For example, the “U”-shaped patterns observed in walleye in boreal reservoirs sites could be explained by the avoidance of some inshore sites shortly after impoundment. Increased water transparency and decreased temperature in those sites may have forced the walleye to redistribute. To isolate the effects of external abiotic forces and climate from the effects of impoundment on fish population dynamics and abundance, we recommend sampling environmental data in reservoirs and in reference sites over time. Clearly very few studies collected comparable fish time series on adjacent reference sites (Figure 3f), and it was difficult to locate companion environmental time series or behavioural data. Biotic factors such as trophic interactions (*e.g*. food, competition and predation; DesLandes *et al*. 1995; Nunn *et al*. 2003; Probst *et al*. 2009) could also explain the variability in patterns observed in our synthesis. In boreal reservoirs, the significant decline in suckers could potentially be explained by increased competition with the whitefish at younger stages (both benthivorous species) and to increased predation by the northern pike (Cook and Bergersen 1988; DesLandes *et al*. 1995). The “U”-shaped pattern observed in whitefish could results from interaction with northern pike. Highest abundance of northern pike in the first years after impoundment coincided with the lowest abundance of whitefish in La Grande 2 and Opinaca reservoirs and the increased of whitefish (right side of the “U”-shaped pattern) coincided with the decline of northern pike (trophic depression).

### No consistent patterns in α- and β-diversity

Our synthesis indicated that species richness, diversity and evenness in reservoirs were comparable to those calculated for reference lakes but we observed an overall higher richness, diversity and evenness in reservoirs and impounded streams compared to un-impounded ones. We also showed that species richness, diversity and evenness do not show consistent directional trends in reservoirs through time and that Jaccard’s analysis failed to find a consistent species turnover pattern across reservoirs.

How can we reconcile our findings in light of the suggested regional homogenization and convergence of fish communities in reservoirs and impounded systems literature? First, it is totally possible that fish faunas converge and become more similar among impounded ecosystems at a regional scale without necessarily changing richness, diversity or evenness within reservoirs. However, we would expect that the process of homogenization across reservoirs would lead to faunal communities becoming more dissimilar over time relative to their pre-impoundment state. Using a Jaccard’ dissimilarity analysis over time on eight reservoirs, we failed to find a consistent species turnover pattern across reservoirs (Appendix D). Some reservoirs became more dissimilar but some became more similar resulting in an overall non-consistent trend. We recognize that the degree of homogenization operating across reservoirs may depend on whether a given species assemblage is constrained by new local habitat characteristics found in reservoirs that could lead to species extirpation, by biotic interactions as well as the introduction and establishment of competitive non-native species. Rahel (2002) reported that an increase in biodiversity can be observed in impoundments as a result of non-native species introduction that outpaced the extirpation of native ones. Non-native species establishment may be facilitated by the lentic habitats created upstream of the dam and by the damped variability in streamflow downstream of the dam (Johnson *et al*. 2008; Clavero and Hermoso 2010). The presence of impoundments may act synergistically with propagule pressure to homogenize species assemblage on a regional scale (Johnson *et al*. 2008) but impoundment on its own is not likely to be the causal factor of fish fauna homogenization. In our synthesis, many reservoirs in the boreal region are characterized by both very low human population density and low propagule pressure, and in which no non-native species have been observed and no native species disappeared (DesLandes *et al*. 1995; Sutela and Vehanen 2008; GENIVAR 2012). Thus, reservoirs in boreal regions might actually be better systems to observe the effects of impoundment in isolation from potentially conflated effects like species invasions on fish fauna assemblages, because they are less affected by the propagule pressure.

The high richness and diversity found in reservoirs might be due to the unique and intermediate position occupied by reservoirs along the “river-lake continuum”. Reservoirs combine features of both river and lake environments, creating a very diverse set of habitats along the reservoir. Vertical gradients of environmental factors in a lentic zone (*i.e*. lake-like ecosystem) are usually found upstream of the dam and they gradually transformed into horizontal gradients in the fluvial zone (*i.e*. river-like ecosystem). The trophic state (*e.g*., turbidity, phosphorus levels, chlorophyll *a*, phytoplankton productivity, dissolved oxygen depletion) may shift from more eutrophic condition closer to the dam to oligotrophic conditions further up in the transitional zone (Kimmel and Groeger 1984; Siler *et al*. 1986; McKenna *et al*. 2005; Okada *et al*. 2005). These gradients should strongly influence fish habitat use, feeding and growth rate resulting in a more diverse fish community (Irz *et al*. 2006). In addition to this mix of vertical and horizontal gradients that should support higher biodiversity in reservoirs, water level fluctuations in reservoirs, if not extreme, might also promote biodiversity through some “diversity-disturbance” relationships (Connell 1978).

Evaluations of faunal communities in reservoirs across the landscape clearly need to have an appropriate reference point, but the answer to the following question is always not self-evident: “What is a reference ecosystem for a reservoir?” Comparing reservoirs to only reference lakes or only rivers might be inadequate to evaluate if fish communities differ from a reference ecosystem because reservoirs are neither a lake nor a river. Irz *et al*. (2006) compared reservoir fish communities with those in rivers and lakes and found that reservoir communities are more similar to lake communities (Jaccard index = 0.10) than river communities (Jaccard index = 0.38; lakes vs. rivers = Jaccard index = 0.43), but this is only one study. Future studies should look at differences in zones along the river-lake continuum in reservoirs, using distance from the dam as an example, to improve our understanding of the spatial and temporal heterogeneity in reservoirs, which in turn could assist in the management reservoir resources (Kimmel and Groeger 1984; Siler *et al*. 1986; Říha *et al*. 2009). We must improve our understanding of how habitat and biological processes are organized in different zones of the reservoirs, how water levels fluctuation impact habitat and biological processes and how this can be translated into fish productivity over time.

The size of ecosystems and their geographic location are other factors that are well known to influence species diversity in lakes (Matuszek and Beggs 1988; Post *et al*. 2000; Samarasin *et al*. 2014) and may have contributed to the variation observed across studies. Indeed, boreal reservoirs used in this synthesis were on average 158 times bigger than adjacent reference lakes (mean ± SD = 158 ± 188 times bigger; range = 10 to 426 times bigger). Therefore, one can expected a higher richness in reservoir relative to adjacent reference lakes just based on their respective size. Based on formulas proposed by Matuszek and Beggs (1988) which control for ecosystem size and by Samarasin *et al*. (2014), controlling for size and location, we would expected 1.9 times and 2.1 times more fish in reservoirs compared to reference lakes, respectively. This suggests that the observed richness, when controlling for size and location, is lower in reservoirs. By applying these formulas to one of our study location, we would estimate that fish richness in the Sainte-Marguerite 3 reservoir (located in North East Québec at 50°57′15.59″N, 66°54′13.18″W) would be between 16 and 33 species. However, this is clearly not possible as there is a potential of only 8-13 species in the region (Chu *et al*. 2003). Many other co-varying variables (*e.g*. pH, dissolved organic carbon, altitude; Matuszek and Beggs (1988)) can also determine the number of species present in a given aquatic ecosystem. Overall, the absence of a directional pattern in richness when comparing reservoirs and reference lakes has the potential to be confounded by numerous factors and thus should be interpreted with caution.

#### Which species generally benefited or are adversely affected by impoundment?

There is an overall consensus in the literature that fish assemblages generally changed in reservoirs relative to pre-impoundment conditions. However, the direction of the change and the species that benefited from, or were adversely affected by impoundment, vary considerably among studies and ecozones. Studies found: 1) an increase in non-native species and a decrease in native species in reservoirs (Martinez *et al*. 1994; Johnson *et al*. 2008; Gido *et al*. 2009; Clavero and Hermoso 2010); 2) a change from a fluvial species dominated community to a more lentic community (Bonner and Wilde 2000; Taylor *et al*. 2001, 2014; Guenther and Spacie 2006; Franssen and Tobler 2013) or 3) a higher abundance of piscivorous fish species in reservoirs and impounded streams than in references ecosystems as a result of introduction and stocking (Quist *et al*. 2005; Guenther and Spacie 2006). We decided to not present data on changes in fish assemblage in reservoirs over time because very different strategies to quantify such a change have been used (*e.g*. ordination techniques such as non-metric multidimensional scaling, species turnover and dissimilarity index, change in proportion of certain guilds and so on). Our analyses comparing the proportion of benthivorous and littoral fish in reservoirs and reference lakes did not turn up significant differences in most cases, but it is possible that this categorization of fish assemblages is too coarse to detect more subtle community changes.

The mechanisms by which fish assemblages change in reservoirs and impounded streams are still not clear but several hypotheses have been suggested, with variable empirical support and often limited to species of commercial and recreational fisheries interests (*i.e*. salmonids, piscivores). Among potential mechanisms, the degree of alteration in the hydrological regime (*i.e*. a change in magnitude, frequency, duration and timing of discharge and water levels; Cohen and Radomski 1993; Gido *et al*. 2000, 2009; Quinn and Kwak 2003; Sutela and Vehanen 2008; Renöfalt *et al*. 2010; Taylor *et al*. 2014), the modification of the riverscape connectivity (Gehrke *et al*. 2002; Guenther and Spacie 2006; Franssen and Tobler 2013) and fish stocking and the deliberate and/or accidental introduction of species in reservoirs (Martinez *et al*. 1994; Rahel 2000; Taylor *et al*. 2001; Quinn and Kwak 2003; Gido *et al*. 2009), have all been suggested to explain the distinct fish assemblage in impounded ecosystems. Moreover, dominant mechanism can change over time (*i.e*. filling phase, shortly after or many years after impoundment) and space (*i.e*. zones of the reservoirs, ecoregions and longitude). The effects of the different mechanisms can also be additive or synergistic, and their effects might strongly depend on reservoir and watershed characteristics and reservoir management.

The alteration of the hydrological regime can affect fish communities by two main processes. First, acting at larger temporal and spatial scales, a general change from lotic to lentic habitats upstream of the dam and a general decrease in discharge downstream of the dam adversely affected fluvial specialist and large-river species adapted to naturally more turbid and fluctuating flow and favored generalist and facultative fluvial species (Winston *et al*. 1991; Bonner and Wilde 2000; Franssen and Tobler 2013; Taylor *et al*. 2014). Second, acting at smaller temporal and spatial scales, water level, discharge fluctuations, and winter drawdown can modify fish assemblages by affecting their feeding, growth and reproduction, markedly for those that depend on the littoral zone. Negative effects can be direct such as the desiccation or freezing of eggs and larvae in the littoral zone, reduced availability of spawning substrate and adequate discharge conditions, reduced growth of juveniles (June 1970; Strange, Fudge & Bodaly 1991; Gafny, Gasith & Goren 1992; Kahl *et al*. 2008; Probst *et al*. 2009) and/or indirect such as alterations in littoral habitat and prey availability and quality (Moyer *et al*. 1995; Paller 1997; Furey, Nordin & Mazumder 2006; Aroviita & Hämäläinen 2008; Zohary & Ostrovsky 2011; Stoll 2013; Sutela *et al*. 2013). For example, Zweiacker, Summerfelt & Johnson (1972) observed a decline in first year growth in largemouth bass in years with important drawdown as a result of its effect on invertebrates. On the other hand, drawdown can favor piscivores by concentrating prey fish, which in turn can improved recreational fishing success in some reservoirs (Hulsey 1956; Ploskey 1986; Nordhaus 1989; Sutela and Vehanen 2008). Increased feeding activity and growth of young and adult piscivores after drawdown has been reported for northern pike, smallmouth bass (Heisey *et al*. 1980), largemouth bass (Heman *et al*. 1969; Zweiacker *et al*. 1972; Herrington *et al*. 2005), white crappie, and flathead catfish (Johnson and Andrews 1973). Water level fluctuations and drawdown can also impact spawning and recruitment of species using the littoral zone (*i.e*. slow–flowing, flooded vegetation dependent and nest spawners; Gasaway 1970; Walburg 1977; Nelson & Walburg 1977; Benson 1980). The modification of the riverscape connectivity by dams (Dynesius and Nilsson 1994; Fullerton *et al*. 2010) can also alter fish assemblages by limiting the movement of migratory species or by facilitating invasions. Populations isolated in upstream areas by dams can be subject to extirpation when reproductive failure or high mortality cannot be counterbalanced by recolonization from downstream sources (Winston *et al*. 1991) or when adverse water quality, altered thermal and hydrologic regimes adversely affect population dynamics (Cushman 1985). On the other hand, some authors have observed increased colonization of non-native species in impounded streams (Havel *et al*. 2005; Johnson *et al*. 2008). Facultative riverine species and fishes introduced to reservoirs can spread downstream past dams and upstream into unimpounded rivers (Winston *et al*. 1991). Interestingly, dams can also act as important barriers to dispersal for exotic invasive and non-native species. In the Laurentian Great Lakes, impoundments have prevented the spread and upstream migration of the exotic sea lamprey in tributary streams (Dodd *et al*. 2003).

Reservoirs are frequently larger and more accessible to humans than are natural lakes, attracting significant recreational fisheries. Because of this, many reservoirs have been subject to intense fish stocking and species introduction (*i.e*. high propagule pressure), mainly for piscivores and sport/game fish species. In addition to a higher susceptibility to propagule pressure, reservoirs are particularly vulnerable to successful establishment of non-native species because they are younger in age and more “disturbed” than natural lakes because they experience a higher rate of fluctuations in water levels, temperature, sedimentation and nutrient content (Thornton *et al*. 1990; Pringle *et al*. 2000; Davis 2003; Didham *et al*. 2007), which can modify predator–prey interactions. Several studies have found an increase in non-native species after impoundment, very often dominated by piscivorous species that were not present or abundant before impoundment (Martinez *et al*. 1994; Quist *et al*. 2005; Guenther and Spacie 2006; Johnson *et al*. 2008; Gido *et al*. 2009; Clavero and Hermoso 2010; Franssen and Tobler 2013). When introduced, they compete with and prey on native species (Li et al. 1987, Minckley et al. 1991). Basses are well known to homogenize fish assemblages by eliminating small-bodied prey species (Jackson 2002) and are very often introduced in temperate reservoirs. Quist *et al*. (2005) found that the fish assemblage switched from a catostomids and cyprinids (*i.e*. river specialists) dominated system prior to impoundment to an exotic species dominated assemblage, mainly piscivores (*e.g*., smallmouth bass, walleye, yellow perch and brown trout), from reaches upstream and downstream of the reservoir.

Time needed to detect a change in fish community is also highly variable among studies and can be as quick and dramatic as 5 years after impoundment (Martinez *et al*. 1994) to more than 30 after impoundment (Quinn and Kwak 2003). Some community states or phases (*i.e*. dominant species in the community) can also be transient with reservoir aging. For instance, Říha *et al*. (2009) and Kubečka (1993) documented a five phase succession in fish species with European reservoirs aging. Fish assemblage changed successively from a riverine species phase (before impoundment and shortly after), to a pike phase (*Esox lucius*), a perch phase (*Perca fluviatilis*), a rapid and transient perch-cyprinid phase to a final and highly stable cyprinid dominated phase. The time needed to detect a significant change in fish assemblage will very likely depend on the dominant mechanisms affecting the community in reservoirs. The number of processes affecting community structure and assemblage, such as competitive interactions or colonisation events, may not have operated long enough to generate detectable community patterns in some reservoirs. If the dominant mechanism is a modification of migration patterns and connectivity, the effect on fish community might be detectable quickly with the extirpation of migratory species. On the other hand, if the dominant mechanism impacting some fish species is water level fluctuation, the effect on fish assemblage may take many years to be detectable.

### Concluding remarks and recommendations

Our quantitative synthesis first highlights that fish recruitment and adult abundance responded quickly and positively to impoundment, but the surge was transient. The direction and magnitude of fish responses to impoundment varied greatly among studies, species, and even among sites for the same species. These findings suggest that monitoring fish populations and community in reservoirs over a couple of years after impoundment is insufficient to assess the potential for fisheries given that many time series were not at equilibrium 15 to 20 years after impoundment. Second, by comparing reservoirs and reference ecosystems distributed from the boreal to the temperate ecozones, we did not find strong empirical evidence for a consistent decrease in species richness, diversity and evenness following impoundment. We also did not find strong evidence for a change in diversity over time or a consistent pattern in species turnover rate following impoundment from a network of reservoirs. These findings suggest that the homogenizing effect of impoundment on fish community previously observed at a global (Rahel 2000, 2002; Poff *et al*. 2007; Liermann *et al*. 2012) and more regional scale (Gido *et al*. 2009; Clavero and Hermoso 2010) might vary from one region to the other (*e.g*., not significant or detectable in boreal ecoregions) and might be the result of several synergistic factors that are confounded with impoundment (*e.g*. propagule pressure).

Monitoring fish for a longer period after impoundment, having reliable data before impoundment and during filling, collecting data on adjacent reference sites (lakes and rivers/streams; BACI design), and avoiding study bias will enhance our abilities going forward to make stronger conclusions about the effect of impoundment and dams on fish populations and communities and will reduce the risk of inappropriate management actions due to some false impressions of very productive fisheries during the surge (Ploskey 1986; Quinn and Kwak 2003). Temporal comparisons in this synthesis clearly demonstrate that short time series resulted in a very different interpretation than the longer time series on the same species in the same reservoir (See Figure 4c and d). Some consequences of impoundment on fish community are intuitive and can be easy to detect (*e.g*., extirpation of migratory species by a dam) whereas others are much more difficult to detect, particularly due to environmental stochasticity and reservoir management (water level fluctuations and mitigations. To avoid this confusion, reference sites (rivers and lakes) are needed (Penczak *et al*. 1998; Głowacki and Penczak 2000).

To statistically isolate mechanisms affecting fish population and community in reservoirs, we need more experimental and field studies at the ecosystem level and habitat variables have to be monitored as well. Fish assemblages change after impoundment but habitats change as well because most reservoirs age more faster that natural lakes (Kimmel and Groeger 1986). Benson (1980) suggested that fish population data collected prior to when reservoir shores have reached a reasonable degree of stability do not provide a reliable estimate of the final species composition in a reservoir. To do so, Management Strategy Evaluation (*i.e*. general framework aimed at designing and testing management procedures) or adaptive management studies would be particularly valuable, where water levels can be manipulate with the collaboration of managers to suit hypotheses and questions of researchers. Facilities such as the Experimental Lakes Area (ELA; 58 small experimental lakes and their watersheds in Northwestern Ontario, Canada) can also be very useful to isolate mechanisms. For example, the water level was lowered during three successive winters in lake 226 from the ELA to evaluate changes in an unexploited lake whitefish population (Mills *et al*. 1999).

Given that this analysis is a data synthesis, we should always keep in mind that the potential for biases at the level of individual studies. For example, one could argue that studies funded by industrial sources might be biased in an opposite direction to those published in peer-reviewed journals (i.e. economic interests might benefit from a “no effect” model whereas academic outlets might be reluctant to publish no effect models). Herein, we have incorporated both types of studies, with the intent of providing a comprehensive and a balanced perspective. To facilitate future syntheses in this area, a greater effort has to be made to use a similar terminology across studies and to compare similar hydrological variables so that all studies can be synthesized to their fullest potential. As suggested by Kimmel & Groeger (1986), an integrated and interdisciplinary research effort of the effects of impoundment and water levels fluctuations on aquatic ecosystems, involving managers and industrial partners, is desirable for the future of reservoir water quality, fisheries, aquatic ecosystem services and integrity.

## Acknowledgements

This work was supported by a MITACS Accelerate grant to IGE, CS and CN, and a MITACS Elevate scholarship to KT with Hydro-Québec as an industrial partner. We thank C. Turpin (Hydro-Québec), A. Tremblay (Hydro-Québec) and P. Johnston (Hydro-Québec) for helpful suggestions and comments on an early draft of this manuscript.

